# Non-canonical aberrant DNA hypermethylation in glioma

**DOI:** 10.1101/193789

**Authors:** Agustin F. Fernandez, Gustavo F. Bayón, Marta I. Sierra, Rocio G. Urdinguio, Estela G. Toraño, Maria García, Antonella Carella, Virginia Lopez, Pablo Santamarina, Thalia Belmonte, Juan Ramon Tejedor, Isabel Cobo, Pablo Menendez, Cristina Mangas, Cecilia Ferrero, Luís Rodrigo, Aurora Astudillo, Ignacio Ortea, Sergio Cueto Díaz, Pablo Rodríguez-Gonzalez, J. Ignacio García Alonso, Manuela Mollejo, Bárbara Meléndez, Gemma Dominguez, Felix Bonilla, Mario F. Fraga

## Abstract

Aberrant DNA hypermethylation is a hallmark of cancer although the underlying molecular mechanisms are still poorly understood. To study the possible role of 5-hydroxymethylcytosine (5hmC) in this process we analyzed the global and locus-specific genome-wide levels of 5hmC in primary samples from 54 gliomas and 72 colorectal cancer patients. Levels of 5hmC in colorectal cancer were very low and no consistent changes were detected between control tissues and tumors. As expected, levels of 5hmC in non-tumoral brain samples were high and significantly reduced at the 49,601 CpG sites in gliomas. Strikingly, hypo-hydroxymethylation at 4,627 (9.3%) of these CpG sites was associated with aberrant DNA hypermethylation. The DNA regions containing these CpG sites were enriched in H3K4me2, and presented a different genuine chromatin signature to that characteristic of the genes classically aberrantly hypermethylated in cancer. We conclude that this data identifies a novel 5hmC-dependent non-canonical class of aberrant DNA hypermethylation in glioma.

## Introduction

DNA methylation at the fifth position of cytosine (5mC) has been one of the most studied epigenetic modifications in mammals to date. 5mC is involved in the regulation of multiple physiological and pathological processes, including cancer, and when located at gene promoters, it is usually linked to transcriptional repression.

As distinctive features of tumorigenesis, local DNA hypermethylation and global hypomethylation have been attributed to changes in 5mC levels [10; 11]. However, the discovery a few years ago, of 5-hydroxymethylcytosine (5hmC), a new epigenetic mark resulting from 5mC oxidation, is reshaping our view of the cancer epigenome [29; 47]. This 5mC to 5hmC conversion in mammals is mediated by ten-eleven translocation proteins (TET1, TET2, and TET3), a family of α-ketoglutarate (αKG) and Fe(II)-dependent dioxygenases[21; 47]. Global levels of 5hmC in the genome fluctuate considerably according to tissue type, and are consistently around 10-fold lower than those of 5mC, though it is interesting that the highest levels of both marks are found in brain [13; 16; 24; 32; 37; 44].

Several studies suggest that 5hmC is not only an intermediate of DNA demethylation, but that it also plays a role in cancer biology [23; 33; 41; 46; 55]. In this vein, a broad loss of 5hmC has been reported in different human cancers including melanoma, glioma, breast, colon, gastric, kidney, liver, lung, pancreatic, and prostate cancers [17; 23; 27; 30; 32; 34; 55].

The fact that there are now methods available that distinguish 5mC and 5hmC positions at single-base resolution within the genome prompted us to reassess the role of DNA methylation status in tumorigenesis from a 5hmC perspective. The method used here allowed us to describe global and genome-wide locus-specific 5mC and 5hmC patterns in colon and brain samples, to identify a specific chromatin signature associated with changes of these epigenetic marks in cancer and, most notably, to describe a novel non-canonical type of aberrant DNA hypermethylation in cancer.

## Results

### Global changes of 5mC and 5hmC in cancer

To evaluate the role of 5hmC in the changes of DNA methylation observed in cancer, we first analyzed the levels of 5hmC and 5mC at repetitive DNA in 84 normal and 123 tumor samples obtained from patients with colorectal cancer and glioma. Bisulfite pyrosequencing was used to determine the level of both epigenetic modifications in 4 different types of repeated DNA: the retrotransposons LINE-1 and AluYb8, and the pericentromeric tandem repeats Sat-alpha and NBL-2 [49]. These 4 DNA regions contain most of the genomic methylation and, consequently, global DNA methylation level is highly dependent on their 5mC content [51]. As expected, 5mC levels at most repeated DNA in healthy tissue was high, and was reduced in tumor samples (**Figure 1A**). In contrast, the levels of 5hmC at repeated DNA in healthy tissue was very low, and tumoral tissue showed even lower levels of 5hmC in these DNA regions (**Figure 1B**). However, the differences were very small and, consequently, they cannot explain the global loss of this mark in cancer observed by mass spectrometry [23; 27; 28; 39].

**Figure 1.**
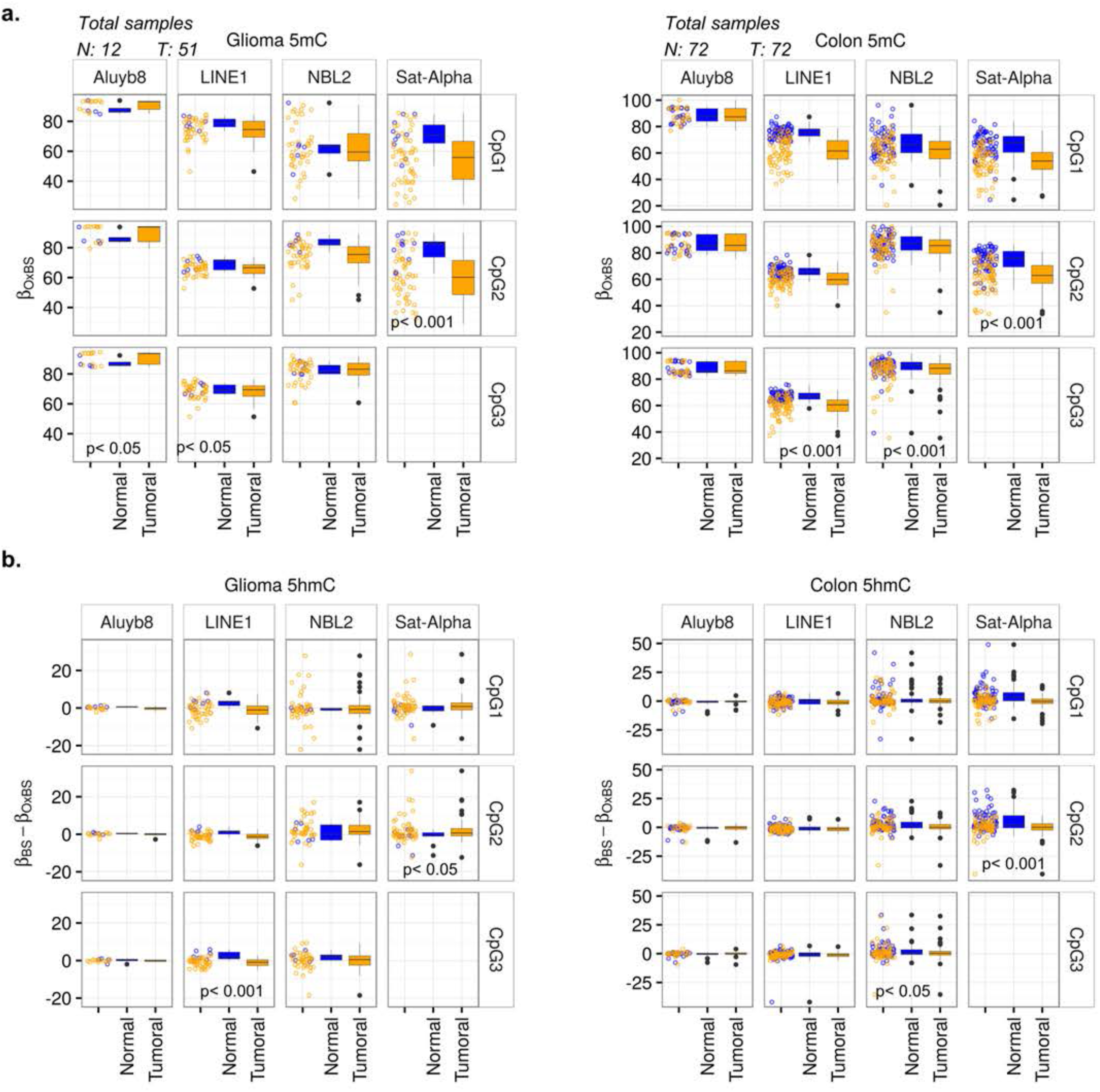
5mC and 5hmC levels at repetitive DNA sequences in glioma and CRC. 5mC (A) and 5hmC (B) values of several repetitive regions (AluYb8, LINE-1, NBL-2, and Sat-alpha) measured by bisulfite pyrosequencing in controls and glioma (left panels) and CRC (right panels). Individual CpG site values for each repeat are displayed. P-values are shown.

### 5mC and 5hmC profiling in colorectal and brain tissue

As changes in 5hmC at repeated DNA were not able to explain the global changes previously observed by mass spectrometry, we hypothesized that these changes primarily occur at single copy sequences. To investigate this possibility in more detail, we first used 450K Infinium methylation arrays to determine the level and genomic distribution of 5mC and 5hmC at 479,423 CpG sites in 11 non-tumoral colorectal samples and 5 healthy brain tissue samples, all from different donors. A preliminary examination of the data revealed that the beta values of the oxidized samples (true 5mC) were much lower than their non-oxidized counterparts (5mC+5hmC) in brain (Wilcoxon rank sum test; p<0.001; W=2.34e13) than in colorectal (Wilcoxon rank sum test; p<0.001; W=5.52e13) tissues (see Materials and Methods), which indicates that, as expected, levels of 5hmC are higher in brain tissue than in the colon (**Figure 2A**). In line with this, we identified 111,633 and 5,089 hydroxymethylated CpG sites (5hmC sites) in brain and colorectal tissue respectively (**Figure 2A, and Supplementary Table 1 and 2**) (see Materials and Methods).

**Figure 2.**
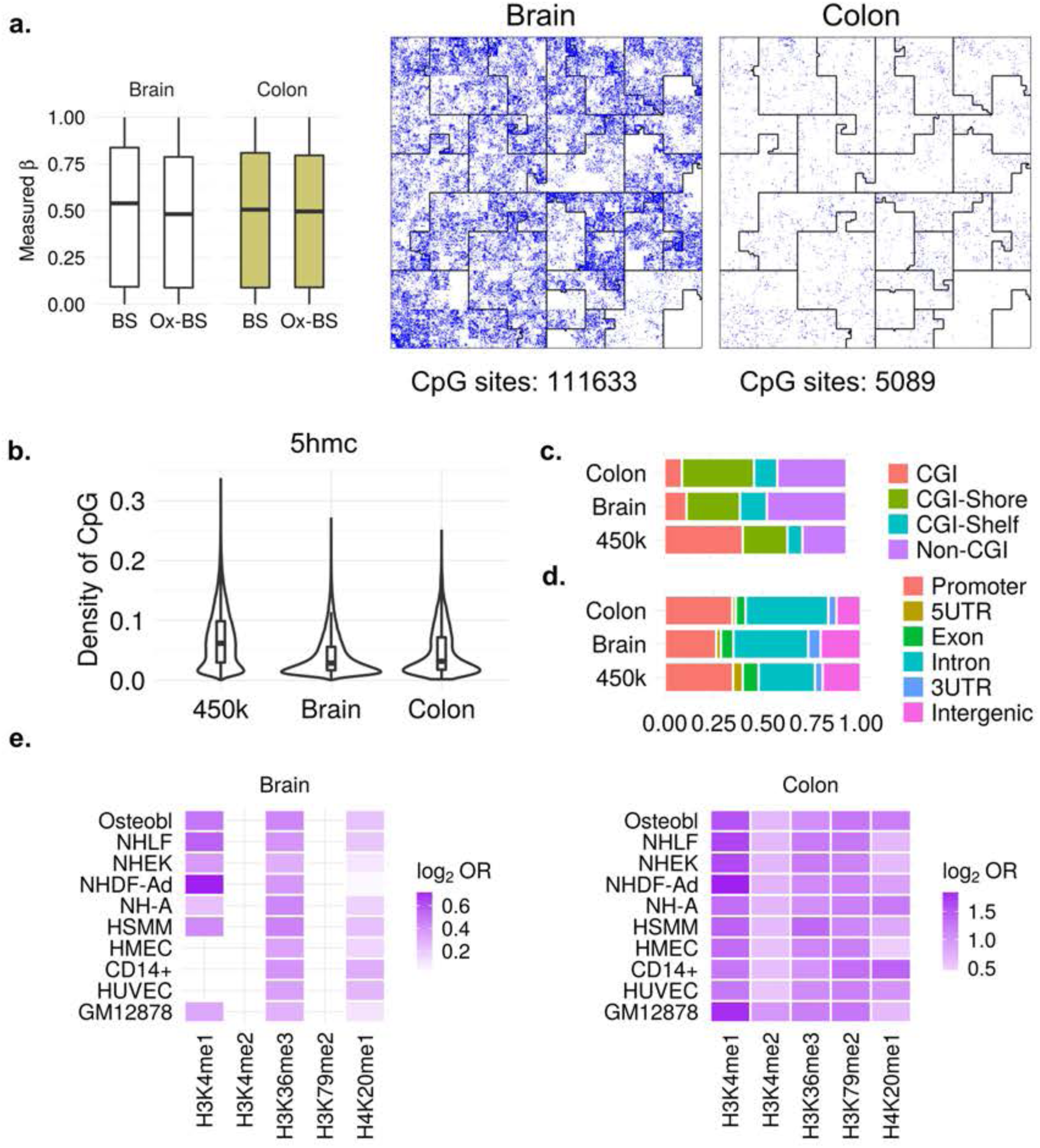
Characterization of DNA 5hmC in normal brain and colon samples. (A) Box plots showing differences between average Beta values of 5mC+5hmC (BS) and true 5mC (OxBS) in both normal brain and colon. On the right are Hilbert curves showing the amount and genomic distribution of 5hmC in brain and colon. (B) Associations between 5hmC and CpG density. (C) Distribution of 5hmC CpG sites relative to CpG island status and compared to the array background (450K). (D) Distribution of 5hmC CpG sites relative to different genomic regions. (E) Heatmaps showing significant enrichment of the 5hmC CpG sites, identified in brain and colon, with different histone marks contained in the UCSC Browser Broad Histone track from the ENCODE project. Color code indicates the significant enrichment based on log2 odds ratio (OR).

The analysis of the genomic distribution of the 5hmC sites showed that, in both colorectal and brain tissue, hydroxymethylation is enriched at the low CpG-density regions interrogated by the array (Wilcoxon non-parametric test; p<0.001, D=-0.29, and p<0.001, D=-0.5, respectively) (**Figure 2B**). Consequently, the 5hmC sites were enriched in non-CpG islands (non-CGI) in both colon and brain (chi-square test; p<0.001; OR=1.93, and p<0.001, OR=3.45, respectively) and infrequent in CGIs (chi-square test; p<0.001, OR=0.14, and p<0.001, OR=0.13) (**Figure 2C**). With respect to genes, 5hmC sites were enriched in introns in both brain and colorectal tissue (chi-square test; p<0.001, OR=1.82, and p<0.001, OR=1.76, respectively), but were less frequent than expected in intergenic regions in colorectal tissue and in gene promoters in brain tissue (chi-square test; p<0.001, OR=0.58, and p<0.001, OR=0.6) (**Figure 2D**). Moreover, hydroxymethylated CpG sites were farther away from centromeres in brain (Wilcoxon non-parametric test, p<0.001, D=0.01) and telomeres in both colorectal tissues and brain (Wilcoxon non-parametric test, p<0.001, D=0.07, and p<0.001, D=0.02, respectively) than the median in terms of other background sites, although the size of these shifts was rather small (**Figure 2-figure supplement 1A**).

To identify possible chromatin marks associated with 5hmC sites in colorectal and brain tissue, we compared these CpG sites with previously published data on a range of histone modifications and chromatin modifiers in 10 different cell types (see Materials and Methods) (**Figure 2E**). This approach identified statistically significant associations (Fisher´s exact test; p<0.05) between the 5hmC sites and the active histone marks H3K4me1, H3K36me3, and H4K20me1, in both colorectal and brain tissue (**Figure 2E**). Interestingly, in colorectal tissue, 5hmC was also enriched in other activating histone posttranslational modifications (PTMs) such as H3K79me2, and H3K4me2 (**Figure2E**). Finally, a similar framework was used to test for the enrichment of our selected probes over the computer-generated chromatin segmentation states from the ENCODE ChromHMM project (see Materials and Methods). In total, fifteen states were used to segment the genome, and these were then grouped and colored to highlight predicted functional elements. This approach showed that the hmC sites were significantly enriched in states associated with enhancers and transcription in both colorectal and brain tissue (Fisher´s exact test; p<0.05) (**Figure 2-figure supplement 1B**).

### Locus-specific alterations of 5hmC in colorectal cancer (CRC) and glioma

To identify differentially hydroxymethylated CpG sites (d5hmC) at single copy sequences in cancer, we used 450K methylation arrays to analyze 11 additional colorectal tumors and 9 primary tumors obtained from patients with glioma (see Materials and Methods). A total of 49,601 CpG sites that were hypohydroxymethylated were identified in gliomas, but almost no hyper-hydroxymethylated sites were found (see Materials and Methods) (**Figure 3A and Supplementary Table 3**). In contrast, no significant methylation changes were found in colorectal tumors (**Figure 3A**) and thus subsequent stages of the study focused on glioma alone. Hierarchical clustering using the differentially hydroxymethylated CpG sites showed the correct classification of normal and tumor samples (**Figure 3B**). The analysis of the genomic distribution of the hypo-hydroxymethylated CpG sites in gliomas showed an enrichment at low CpG density regions (Wilcoxon rank sum test, p<0.001, D=-0.41), and consequently at non-CpG islands (chi-squared test, p<0.001, OR=2.53) (**Figure 3C**). With respect to gene location, hypo-hydroxymethylation was more frequent in introns (chi-squared test, p<0.001, OR=1.77) (**Figure 3C**).

**Figure 3.**
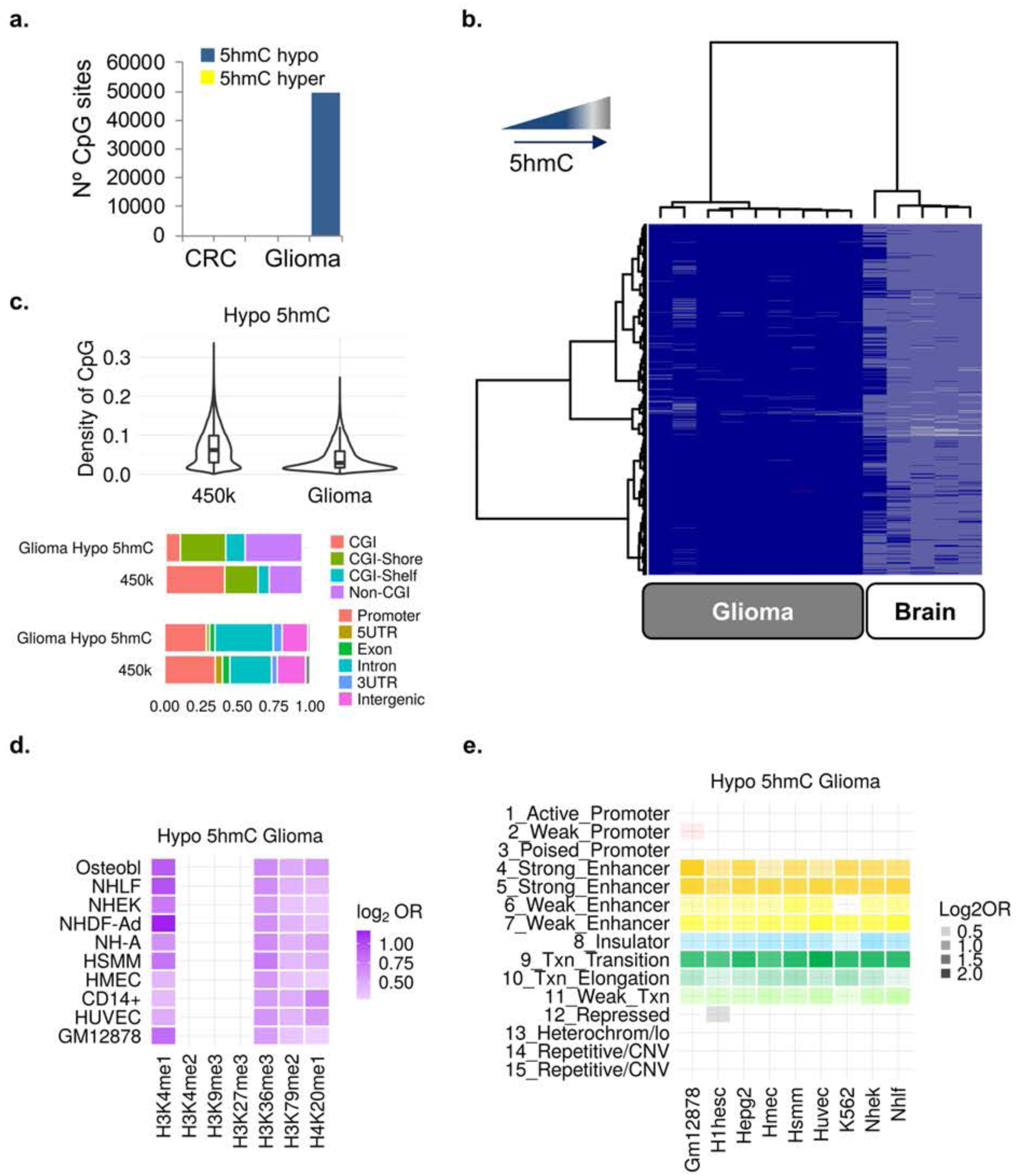
Alterations of 5hmC in CRC and glioma. (A) Bar plot showing the number of dh5mC sites in CRC and glioma. (B) Unsupervised hierarchical clustering and heatmap including CpG sites with 5hmC loss in glioma. (C) Associations between 5hmC loss in glioma and density of CpGs (upper panel), CpG island status (middle panel), and different genomic regions (lower panel). (D) Heatmaps showing significant enrichment of hypo 5hmC CpGs identified in glioma with different histone marks contained in the UCSC Browser Broad Histone track from the ENCODE project. (E) Heatmaps showing significant enrichment of hypo 5hmC CpGs in gliomas with fifteen “chromatin states” generated by a Hidden Markov Model (HMM) (right panel). Color codes indicate the significant enrichment based on log2 odds ratio (OR).

To identify possible chromatin signatures associated with DNA hypo-hydroxymethylation in gliomas, we compared our list of hypo-hydroxymethylated CpG sites with previously published data on a range of histone modifications and chromatin modifiers in 11 different cell types (see Materials and Methods) (**Figure 3D**). Interestingly, this approach showed an enrichment of hypo-hydroxymethylation at chromatin regions marked with the activating histone PTMs H3K4me1, H3K36me3, H4K20me1 and H3K79me2 (Fisher's exact test, p<0.05) (**Figure 3D**), but not with the repressive histone modification H3K27me3, which has been previously shown to be associated with aberrant DNA hypermethylation in cancer [38; 52] (**Figure 3D**). A similar framework was used to test for the enrichment of our selected probes over the computer-generated chromatin segmentation states from the ENCODE ChromHMM project. Using this approach, we found that hypohydroxymethylated CpG sites were significantly associated with transcription regulation and enhancers (Fisher´s exact test; p < 0.05) (**Figure 3E**).

### DNA hypo-hydroxymethylation identifies a novel type of non-canonical aberrant DNA hyper-methylation in glioma

To study the relationship between changes in 5mC and 5hmC in glioma, we first identified aberrantly methylated CpG (d5mC) sites. The comparison of the methylation data between tumoral and control samples (see Materials and Methods) identified 2,727 hypo- and 12,050 hyper-methylated CpG sites in gliomas (**Supplementary Tables 4 and 5**). Next, we compared these d5mC sites with the previously identified hypo-hydroxymethylated CpG sites (**Figure 3A, Supplementary Table 3**). Interestingly, this approach showed that 4,627 (38.4%) of the CpG sites aberrantly hypermethylated in gliomas also lose 5hmC (**Figure 4A, Supplementary Table 6**).

**Figure 4.**
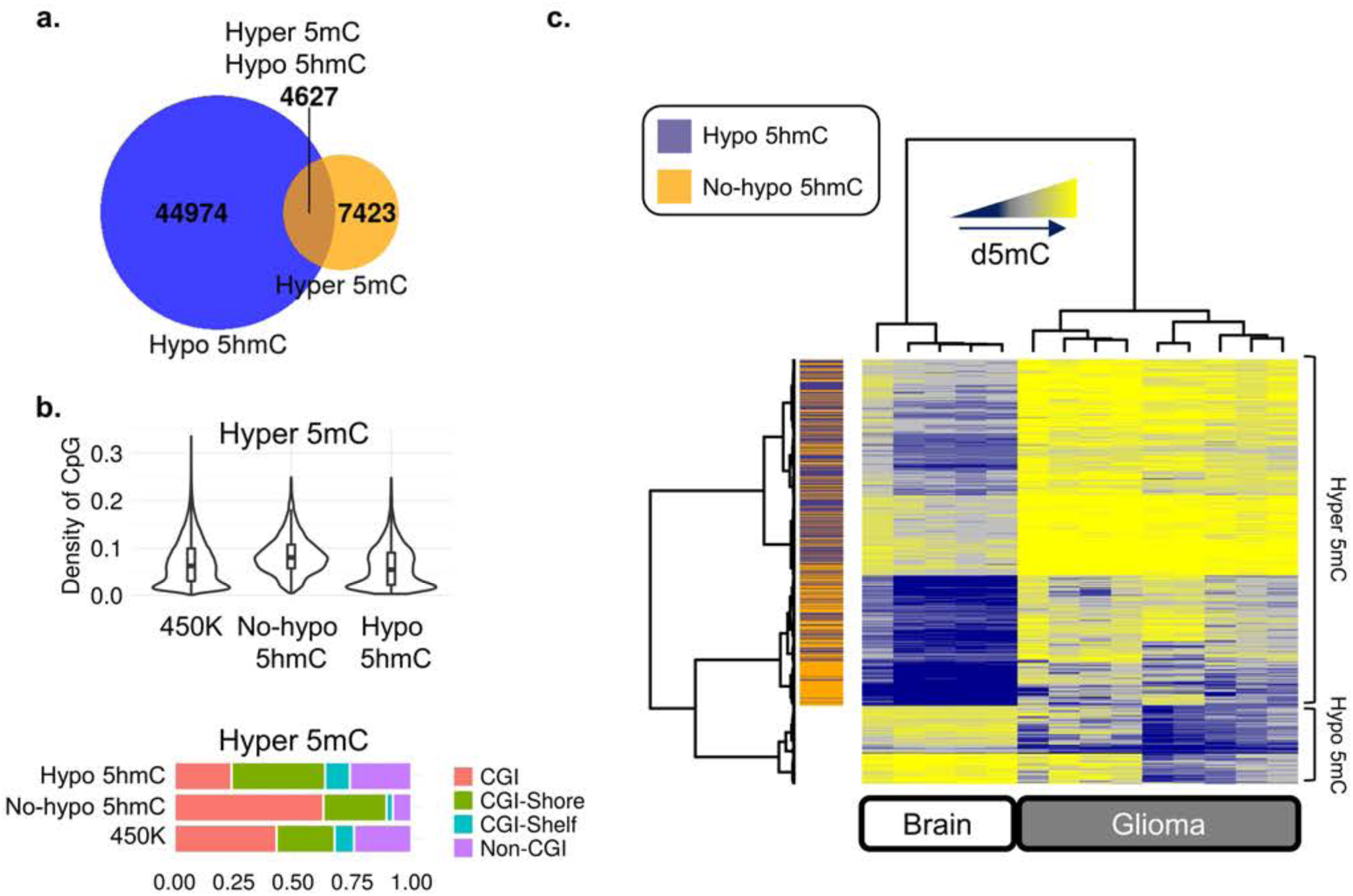
Relationships between changes in 5mc and 5hmc in glioma. (A) Euler diagram illustrating overlap of CpGs that lose 5hmC (hypo 5hmC) and gain 5mC (hyper 5mC) in glioma. (B) Associations between hypermethylated CpG sites that lose (or not) 5hmC and CpG density and CpG island status, compared to the array background (450K). (C) Unsupervised hierarchical clustering and heatmap including CpG sites with 5mC changes (hyper- and hypomethylation) in glioma. Hypo-(purple) and non-hypo (orange) 5hmC overlapped CpGs are indicated by colored lines on the annexed track. Average beta methylation values are displayed from 0 (blue) to 1 (yellow).

To investigate, at a functional genomic level, the characteristics of these two classes of aberrantly hypermethylated CpG sites in gliomas we first analyzed their genomic distribution in relation to density of CpG sites and we found that the hypermethylated CpG sites that lose 5hmC (hyper5mC-hypo5hmC) were enriched in low density CpG regions (Wilcoxon rank sum test, p<0.001, D=-0.11) as compared with the hypermethylated CpG sites that showed no changes in 5hmC (hyper5mC) (Wilcoxon rank sum test, p<0.001, D=-0.23) (**Figure 4B, Supplementary Tables 6 and 7**). In line with this, hyper5mC-hypo5hmC sites were strongly depleted from CGIs (chi-squared test, p<0.001, OR=0.42) (**Figure 4B**). Hierarchical clustering using the differentially methylated CpG sites showed that the hyper5mC-hypo5hmC sites were slightly more methylated in control brain samples than the hyper5mC sites, and that they were more uniformly hypermethylated in glioma (**Figure 4C**). To further corroborate our results, we took advantage of recently published data on the whole-genome bisulfite sequencing (WGBS) in glioma [42]. We found that, in addition to a large percentage of CpGs (n: 4,051; 88%) showing the same patterns of change as in our methylation arrays, the WGBS analysis identified more than 10^6^ new hyper5mC-hypo5hmC sites, thus confirming that this is a frequent event in glioma (**Figure 4-figure supplement 1**).

Next, to identify possible chromatin signatures associated with the two classes of aberrantly hypermethylated CpG sites in gliomas, we compared our data with previously published data on a range of histone modifications and chromatin modifiers in 11 different cell types (see Materials and Methods) (**Figure 5A**). This approach confirmed the association between hyper5mC and the repressive histone marks H3K9me3 and H3K27me3 (Fisher's exact test, p<0.05) [36; 38; 52]. The hyper5mC-hypo5hmC sites showed a completely different chromatin signature, with enrichment in the activating histone PTMs H3K4me1, H3K36me3, H3K79me2 and H4K20me1 (Fisher's exact test, p<0.05) (**Figure 5A**). Notably, as compared with the chromatin signature of the whole set of hypo-hydroxymethylated CpGs in glioma, these CpG sites were particularly enriched at the H3K4me2 histone mark (Fisher's exact test, p<0.001, OR in [1.19, 1.78] for all cell lines in the Broad Histone project) (**Figure 5B**).

**Figure 5.**
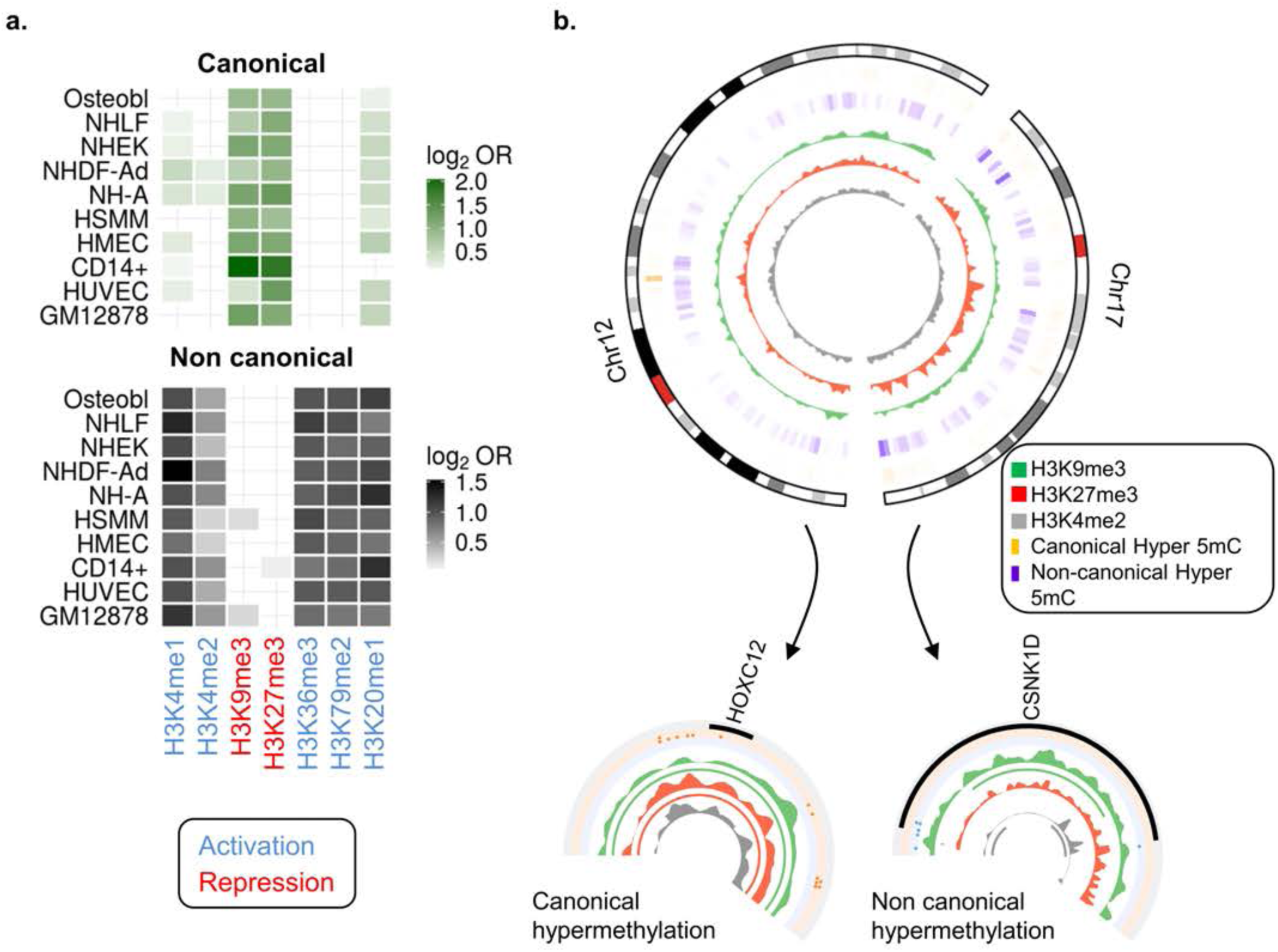
Canonical and non-canonical hypermethylation in glioma. (A) Heatmaps showing significant enrichment of CpG sites in glioma which exclusively gain 5mC (canonical hypermethylation) (upper panel), and both lose 5hmC and gain 5mC (non-canonical hypermethylation) (lower panel), with different histone marks contained in the UCSC Browser Broad Histone track from the ENCODE project. Histone PTMs related to activation and repression are distinguished by colors as indicated in the key. (B) Circular representation of two representative chromosomes (12 and 17), indicating genomic location of canonical (orange) and non-canonical (purple) hypermethylation in glioma. Inner tracks display chromatin marks (H3K9me3, H3K27me3, and H3K4me2), generated for NH-A cells. Two examples of genes showing canonical and non-canonical hypermethylation associated with specific chromatin signatures are displayed below.

These results indicate that the hyper5mC sites behave like the aberrantly hypermethylated canonical CpG sites in cancer, whilst the hyper5mC-hypo5hmC sites represent a novel and functionally different non-canonical type of aberrantly methylated DNA sequence in glioma (**Figure 5A**, **5B**, **Supplementary Tables 6 and 7**). In support of this notion, experiments focused on the computational prediction of functional elements confirmed the enrichment of canonical aberrant hypermethylation in promoters and repressed sequences and revealed a completely different pattern for non-canonical hypermethylation, one which is more closely associated with enhancers and transcriptional regulation (Fisher´s exact test; p < 0.05) (**Figure 5-figure supplement 1**).

### Distinct functional role of canonical and non-canonical aberrant hypermethylation in glioma

To identify possible differences between the functional role of canonical and non-canonical aberrant DNA hypermethylation in glioma we first ascribed CpG sites to specific genes and then used HOMER to carry out gene ontology analyses of each group of genes (see methods). Using this approach, we identified 1,921 genes displaying canonical hypermethylation, 2,042 displaying non-canonical hypermethylation and 938 displaying both types of aberrant hypermethylation (**Figure 6A, Supplementary Tables 8, 9 and 10**). As expected, GO analyses showed an enrichment of development and differentiation processes in canonical genes [6] (**Figure 6A, Supplementary Table 11**). In contrast, non-canonical genes were strongly enriched in cell signaling and protein processing pathways (**Figure 6A, Supplementary Table 12**).

**Figure 6.**
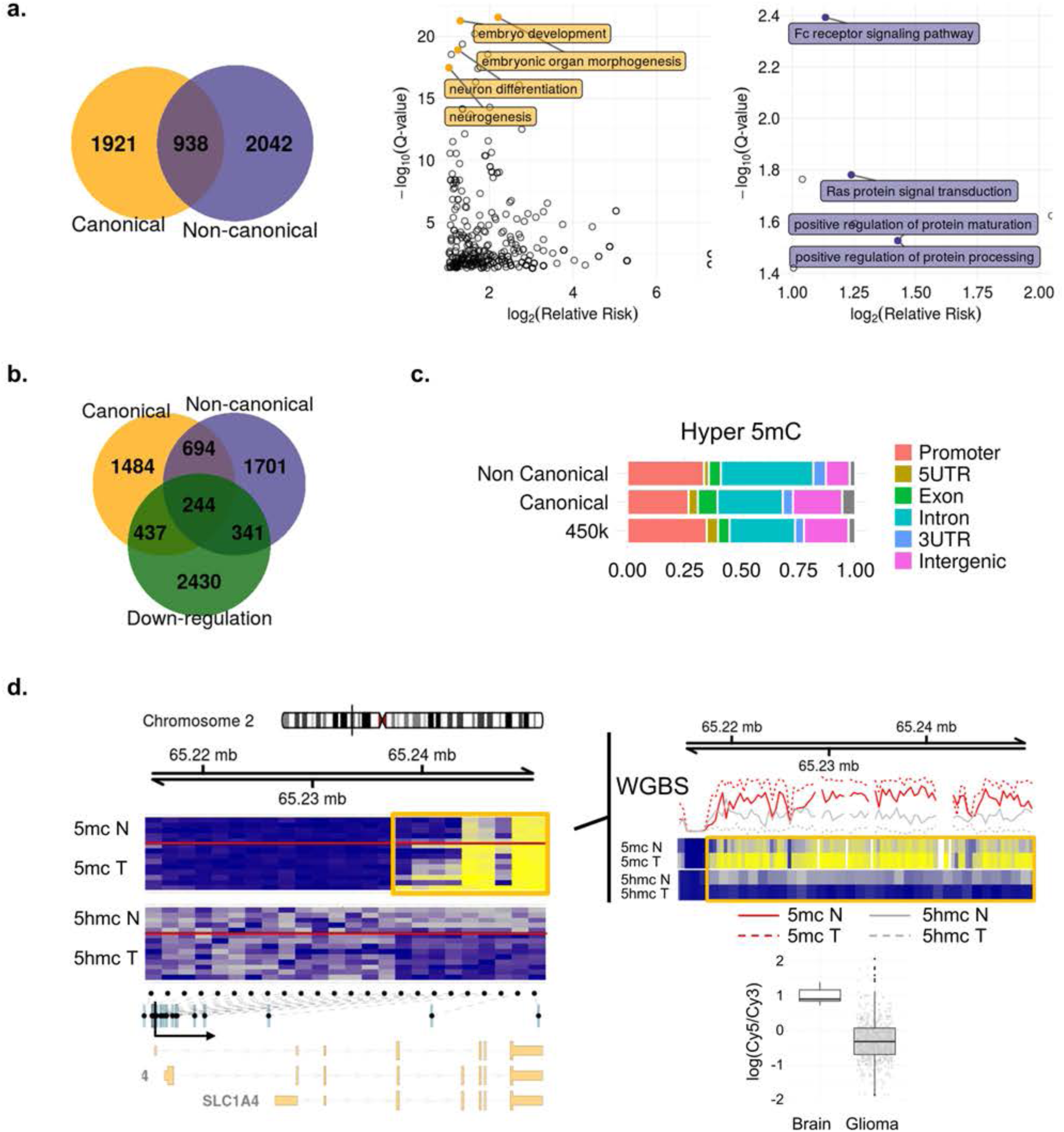
Functional role of canonical and non-canonical hypermethylation in glioma. (A) Euler diagrams showing number of genes associated with canonical hypermethylation, non-canonical hypermethylation, or both. On the right are representative gene ontology terms (Biological process) of genes associated with canonical (orange) and non-canonical (purple) hypermethylation, ranked by Q-value, and enrichment score (relative risk). (B) Euler diagram showing overlap of canonical and non-canonical hypermethylated genes with down-regulation. (C) Associations of canonical and non-canonical hypermethylation in glioma with different genomic regions. (D) Representative example of one gene (*SLC1A4*) showing non-canonical hypermethylation in glioma (orange frame). Organization of the gene, locations of CpGs included in the methylation array (black dots), and transcription start site (TSS) are shown below. 5mC hypermethylation (blue to yellow) and 5hmC loss (gray to blue) in glioma are shown above. Whole genome bisulfite sequencing (WGBS) data [42] including all the CpG sites in the same region are shown on the right. The associated change in gene expression is displayed below.

To further investigate the functional role of canonical and non-canonical hypermethylation in cancer, we compared our methylation data with previously published gene expression data in the same type of tumor (see Materials and Methods). Results showed that 681 (31%) of the canonical and 585 (24%) of the non-canonical aberrantly hypermethylated genes were repressed in gliomas (**Figure 6B**).

Genomic distribution analysis of both types of aberrant hypermethylation confirmed the enrichment of canonical hypermethylation in exons (chi-squared test, p<0.001, OR=1.79 for general exons, OR=2.01 for first exons), while non-canonical hypermethylation was more frequent in introns (chi-squared test, p<0.001, OR=1.7) (**Figure 6C**). The genes frequently downregulated in glioma, *SLC14A* and the *SMAD7*, represent two bona fide examples of this pattern of non-canonical aberrant hypermethylation (**Figure 6D, Figure 6-figure supplement 1**).

Taken as a whole, these results indicate that both types of aberrant hypermethylation have a similar effect on gene expression, but that they affect different types of genes and gene regions.

## Discussion

During recent decades, it has largely been accepted that aberrant genomic DNA methylation is a hallmark of cancer [10; 11] and the best-known DNA methylation alterations in tumors were the aberrant hypermethylation of CpG island promoters, and global DNA hypomethylation. In both cases, the alterations were mostly attributed to changes in the overall content and genomic distribution of 5mC [10; 11].

The vast majority of studies on DNA methylation and cancer have been based on the sodium bisulfite modification of the genomic DNA, a chemical reaction that allows C and 5mC to be distinguished by polymerase chain reaction [19]. However, this approach cannot distinguish between 5mC and 5-hydroxymethylcytosine (5hmC), the latter being a chemical modification of the cytosine first identified in bacteriophages in 1952 [54], and which has recently been found to be quite abundant in specific mammalian tissue [29]. 5hmC is synthesized from 5mC by the Ten-eleven Translocation (Tet) Enzymes, a family of proteins that can also catalyze the successive conversion of 5hmC to 5-formylcytosine and then to 5-carboxylcytosine, both of which can be transformed to unmodified C [40]. Although 5hmC was originally described as simply a demethylation intermediate of C [16; 25], recent data suggest that this may be an epigenetic mark in its own right [3; 22]. Thus, as most previous studies did not distinguish between 5mC and 5hmC, and it appears that DNA hydroxymethylation might play a specific role in cancer, in this work we aimed to re-evaluate changes in DNA methylation in cancer, paying special attention to the specific contribution of 5hmC.

To identify the DNA regions affected by hydroxymethylation changes in cancer, we first focused on four types of repeated DNA (LINE1, Sat α, NBL2 and ALUYB8). Among them, the LINE1 repeat is of particular interest because it contains almost 20% of the genomic 5mC, and it has been proposed to be a surrogate of global DNA methylation [51]. Our results confirmed that tumors lose 5mC at repeated DNA [7]. However, the level of 5hmC at repeated DNA in healthy samples was very low and no significant differences were observed compared to tumors, which indicates that the global DNA hypo-hydroxymethylation previously observed in cancer [23; 27; 28; 39] does not principally occur at repeated DNA. As changes in 5hmC at repeated DNA could not explain the global differences previously observed by mass spectrometry, we decided to study the possible contribution of single copy sequences. Genome-wide profiling of 5mC and 5hmC of healthy tissue identified a 10-fold increase in abundance of CpG sites frequently hydroxymethylated in brain compared to colorectal tissue, providing evidence that the level of this epigenetic mark is highly tissue type-dependent, and also that it is very abundant in the brain [16; 24; 29; 32; 37; 44]. Moreover, 5hmC was enriched in regions with low CpG density and in introns in both colorectal and brain tissue. As the 5hmC is enriched in different genomic regions, these results support the notion that 5hmC is not simply a demethylation intermediate [1]. Interestingly, 5hmC co-localized in regions marked with the activating histone PTM H3K4me1. This histone mark has been previously associated with gene enhancers [20; 48], which suggests that DNA hydroxymethylation might play a role in gene regulation in trans. Moreover, we have recently found an association between H3K4me1 and DNA hypomethylation during aging in stem and differentiated cells [12], which may represent an interesting link between aging and cancer at these genomic regions. Colorectal tumors showed more changes with respect to 5mC than to 5hmC. However, in contrast, glioma presented more changes in 5hmC than in 5mC, suggesting that the dynamics of DNA methylation and hydroxymethylation in cancer is highly tumor-type dependent. Moreover, the great number of hypo-hydroxymethylated single CpG sites in glioma could explain the global differences previously observed by mass spectrometry [23; 27; 28; 39] and suggests that, in contrast to 5mC, most DNA hypo-hydroxymethylation in brain tumors occurs at single copy sequences.

The behavior of 5hmC led us to next identify two types of CpG sites aberrantly hypermethylated in glioma: aberrantly hypermethylated CpG sites that showed no changes in 5hmC; and hypermethylated CpG sites that lose 5hmC. The first of these sites display similar chromatin signatures to previously described genes aberrantly hypermethylated in cancer (i.e. enrichment in the repressive histone marks H3K9me3 and H3K27me3) [36; 38; 52]. In contrast, the second type of aberrantly hypermethylated CpG sites were enriched in the activating histone PTMs H3K4me1, H3K36me3, H3K79me2, H4K20me1 and H3K4me2. As these CpG sites present a genuine chromatin signature which is different to the repressive chromatin signature of the classical genes aberrantly hypermethylated in cancer [36; 38; 52], we conclude that they represent a novel 5hmC-dependent non-canonical class of aberrant DNA hypermethylation in glioma. As this gain in 5mC is inversely correlated with loss of 5hmC, it was not possible to identify this significant alteration in previous studies using the classical sodium bisulfite-based technologies, since they are not able to distinguish between the two chemical modifications.

Aberrant DNA hypermethylation in cancer was discovered more than 30 years ago, but the underlying molecular mechanisms are still poorly understood. For example, it has been proposed that genes enriched in bivalent histone modifications (H3K4me3 and H3K27me3) and polycomb group proteins during embryo development are prone to become aberrantly hypermethylated in cancer [36; 38; 52] but the molecular basis of this is unknown. Our data suggest that tumor cells might in fact acquire aberrant DNA methylation through various different pathways. Moreover, in the case of the non-canonical hypermethylation, the previous loss of 5hmC suggests that aberrant hypermethylation at these DNA regions could be due to an attempt by the cell to reverse or repair the loss of 5hmC at functionally sensible loci. This possibility is supported by the fact that the non-canonical aberrant hypermethylation described here seems to play an important role in gene regulation. Intriguingly, 5hmC at gene promoters has also been proposed to protect from aberrant hypermethylation in colorectal cancer [50]. Thus, although it seems that 5hmC plays an important role in the regulation of the DNA methylation changes in cancer, more research is needed to fully understand its role.

The non-canonical aberrant hypermethylation described here seems to have a similar overall effect on gene expression as classical canonical hypermethylation, although the type of genes and the genomic regions affected are very different. Previous research has shown that the repression of developmental genes affected by canonical aberrant hypermethylation promotes tumorigenesis [6]. However, the possible functional role of disruption of cell signaling and protein processing pathways affected by the non-canonical hypermethylation described in this study remains to be elucidated. Future research is thus needed to address this issue, and to determine whether the two types of aberrant DNA hypermethylation have distinct functional roles in cancer.

## Materials and methods

### Normal samples and primary tumors

The colon and brain samples analyzed in this study were collected at the Hospital Universitario Central de Asturias (HUCA), the Hospital Virgen de la Salud, Toledo, and the Hospital Universitario Puerta de Hierro, Madrid. The samples studied comprised 72 normal colons, 13 normal brains, 72 colorectal primary tumors and 54 glioblastomas. The study was approved by the Clinical Research Ethics Committee and all the individuals involved provided written informed consent.

### Pyrosequencing assays

5mC and 5hmC patterns at repetitive sequences (LINE1, ALUBY8, Sat α and NBL2) were analyzed by pyrosequencing using previously described primers [49]. To calculate 5hmC levels, each sample was analyzed using two methods performed in parallel; an oxidative bisulfite conversion (oxBS) and a bisulfite-only conversion (BS), in accordance with the TrueMethyl^®^ Array Kit User Guide (CEGX, Version 2) with some modifications. Briefly, DNA samples were cleaned using Agencourt AMPure XP (Beckman Coulter) then oxidated with 1 μL of a KRuO4 (Alpha Aeser) solution (375 mM in 0.3 M NaOH), after which bisulfite conversion was performed using EpiTect bisulfite kit (Qiagen^®^).

After PCR amplification of the region of interest in oxBS and BS samples, pyrosequencing was performed using PyroMark Q24 reagents, and vacuum prep workstation, equipment and software (Qiagen^®^). 5hmC levels were obtained when methylation values of oxBS samples (represents true 5mC) were subtracted from their corresponding BS treated pairs (the latter representing 5mC+5hmC).

### Genome-wide DNA methylation analysis with high-density arrays

Microarray-based DNA methylation profiling was performed with the HumanMethylation 450 BeadChip [2]. Oxidative bisulfite (oxBS) and bisulfite-only (BS) conversion was performed using the TrueMethyl^®^ protocol for 450K analysis (Version 1.1, CEGX) following the manufacturer’s recommended procedures. Processed DNA samples were then hybridized to the BeadChip (Illumina), following the Illumina Infinium HD Methylation Protocol. Genotyping services were provided by the Spanish Centro Nacional de Genotipado (CEGEN-ISCIII) (www.cegen.org). DNA methylation data were downloaded from ArrayExpress accession numbers E-MTAB-6003 (brain) and E-MTAB-xxx (colon).

### HumanMethylation450 BeadChip data preprocessing

Raw IDAT files were processed using the R/Bioconductor package minfi [15] (version 1.14.0), implementing the SWAN algorithm [35] to correct for differences in the microarray probe designs. No background correction or control probe normalization was applied. Probes where at least two samples had detection p-values > 0.01, and samples where at least 5500 probes had detection p-values > 0.01 were filtered out. M-values and beta values were computed as the final step in the preprocessing procedure. In line with a previously published methodology [5], M-values were used for the statistical analyses and beta values for effect size thresholding, visualization and report generation.

### Batch effect correction

In order to detect whether there was any batch effect associated with technical factors, the visualization technique of multidimensional scaling (MDS) was employed to highlight any strange interaction affecting the different samples. Where necessary, posterior adjustment of the samples was performed by means of the SVA method [31] implemented in the R/Bioconductor sva package (version 3.14.0).

### Computation of hydroxymethylation levels

Beta values from oxBS samples were subtracted from their corresponding BS treated pairs, generating an artificial dataset representing the level of 5hmC for each probe and sample as per a previously published methodology [45]. One further dataset was created to represent the 5mC levels using beta values from oxBS samples.

### Detection of differentially methylated probes

Differential methylation and hydroxymethylation of an individual probe was determined by a moderated t-test implemented in the R/Bioconductor package limma [43]. A linear model, with methylation or hydroxymethylation levels as response and the sample group (normal/tumoral) as the principal covariate of interest, was then fitted to the methylation or hydroxymethylation data. Surrogate Variables generated using SVA were also included in the model definition, but excluding those found to be correlated to the phenotype of interest. P values were corrected for multiple testing using the Benjamini-Hochberg method for controlling false discovery rate (FDR). An FDR threshold of 0.001 was employed to determine differentially methylated and hydroxymethylated probes. Additionally, these probes were filtered according to their effect size, keeping only those probes with methylation or hydroxymethylation changes between-groups which exceeded the median of all differences for the same comparison. The probes without no significant 5hmC signal on control samples were filtered out from the set of hypo-hydroxymethylated probes in glioma.

### Identification of hydroxymethylated probes

In order to identify those probes representing the regions where the 5hmC mark is located, a differential hydroxymethylation analysis was performed as described previously [14] using a dataset containing both oxBS and BS versions of the control samples. Probes with significant differences in beta values between the BS and oxBS samples were considered to be enriched for the 5hmC mark. An FDR threshold of 0.001 was employed. No filtering on effect size was applied in this case.

### Histone enrichment analysis

In order to analyze the enrichment of histone marks on a subset of probes, we used the information contained in the UCSC Genome Browser Broad Histone track from the ENCODE Project. Histone mark peaks were downloaded for every combination of cell line and antibody. For each track, a 2x2 contingency table was built to represent the partition of the whole set of possible probes in the microarray with respect to the membership of the subset of interest and the overlap between the probes and the histone peaks. A Fisher's exact test was used to determine whether there was significant enrichment of the selected histone mark for the subset of interest. P-values were adjusted for multiple comparisons using the Benjamini-Hochberg method for controlling FDR. A significance level of 0.05 was used to determine whether the given combination of histone mark and cell line presented a significant change in proportion. Additionally, the base-2 logarithm of the Odds Ratio (OR) was used as a measure of effect size.

### Chromatin segment enrichment analysis

Data from the BROAD ChromHMM Project were downloaded from the UCSC Genome Browser site. Each of the tracks comprising this dataset represents a different segmentation generated by a Hidden Markov Model (HMM) using Chip-Seq signals from the Broad Histone Project as inputs. The segmentations were later curated and labelled according to their functional status [8; 9]. In order to detect any significant enrichment in the proportion of probes in a given subset of interest belonging to one functional category, an analysis strategy similar to the one employed for the detection of histone enrichment was performed. In this case, a 2x2 contingency table was built using segments of a given functional status rather than antibodies. A Fisher's exact test was employed, and significant combinations were detected using a FDR threshold of 0.05 (Benjamini-Hochberg procedure). Again, the base-2 logarithm of the OR was used as a measure of effect size.

### Genomic region analysis

The probes in the microarray were assigned to a genomic region according to their position relative to the transcript information extracted from the R/Bioconductor package TxDb.Hsapiens.UCSC.hg19.knownGene (package version 3.1.2). A probe was said to be in a promoter region if it was located in a region up to 2kb upstream of the transcription start site (TSS) of any given transcript. Similarly, a set of mutually exclusive regions were defined inside the transcripts, namely 5UTR, 3UTR, First Exon, Exon and Intron. A probe could only belong to one category, hence if the location of a probe overlapped with two or more regions in different transcripts, it was assigned to the region with a higher level of precedence (i.e. in the order stated above, earlier mention indicates higher precedence). If a probe was not assigned to any of these special regions, it was labelled by default as Intergenic. A contingency table was built for each of the subsets, partitioning the whole set of probes according to membership to a given category and the subset of interest. A Pearson's χ2 test was used to determine whether there was any significant change in proportion between the number of probes marked as belonging to a given region inside and outside the subset of interest. A significance level of 0.05 was employed, and effect size measured by OR.

### CGI status analysis

Similar to the genomic region analysis, probes were labelled according to their relative position to CpG-islands (CGIs), the locations of which were obtained from the R/Bioconductor package FDb.InfiniumMethylation.hg19 (package version 2.2.0). The generation procedure of these CGIs is described by [53], i.e. ‘CpG shores’ were defined as the 2kbp regions flanking a CGI. ‘CpG shelves’ were defined as the 2kbp regions either upstream of or downstream from each CpG shore. Probes not belonging to any of the regions thus far mentioned were assigned to the special category ‘non-CGI’ with each probe being assigned to only one of the categories. A 4x2 contingency table was constructed for each subset of probes in order to study the association between the given subset and the different CGI categories. A χ2 test was used to determine whether any of the categories had a significant association with the given subset. For each of the CGI status levels, a 2x2 contingency table was defined and another χ2 test used to independently evaluate the association of the given subset with each status level, a significance level of 0.05 being employed for all tests. Effect size was reported as the OR for each of the individual tests.

### Analysis of CpG density

For each of the probes in the HumanMethylation450 microarray, CpG density was measured as the number of CG 2-mers present divided by the number which would be theoretically possible in a 2kbp window with the CpG under study at its centre. A Wilcoxon non-parametric test was used to determine if any significant difference existed between the CpG density of each subset of interest and that of the array probes in the background. A significance level of 0.05 was employed for all tests. Effect size was measured using Cliff's Delta (D).

### Gap distance analysis

Distance to both the centromere and telomere was measured for each of the probes in the HumanMethylation450 microarray. In order to find significant differences between the probes within the subset of interest and those in the background, a Wilcoxon non-parametric test was used. Once again, a significance level of 0.05 was employed for all tests, and Cliff's Delta (D) was used as a measure of effect size.

### Microarray background correction

Although it is sometimes referred to as a genome-wide solution, the HumanMethylation450 BeadChip only covers a fraction of the entire genome. In its 27K predecessor, the probes were mainly located at gene promoter regions, while the newer HumanMethylation450 BeadChip additionally includes probes located inside genes and in intergenic regions [4].

The irregular distribution of probes can however lead to unwanted biases when studying whether a selected subset of probes is enriched with respect to any functional or clinical mark. For this reason, here a reference to the background distribution of features was included in all statistical tests performed in order to prevent our conclusions from being driven by the irregular distribution of probes. In qualitative tests (CGI status, genomic region, or histone mark enrichment), the contingency matrix was built to represent the background distribution of the microarray. In quantitative tests (CpG density, distance to centromeres and telomeres) the corresponding metric was compared between the subset of interest and the remaining probes in the microarray. Thus, any significant result would indicate a departure from the fixed background distribution and ignore any bias inherent in the test.

### Gene ontology analysis and annotation

Probe sets were converted to gene sets by using the annotation information from the R/Bioconductor package TxDb.Hsapiens.UCSC.hg19.knownGene (version 3.1.2). A probe was assigned to a gene if the probe was contained within the overlap of all the genomic regions represented by the different transcripts belonging to that gene, or in a 2kbp region upstream of the corresponding TSS. Probes converted this way can be assigned to one or more genes, or to zero (i.e. intergenic probes).

After gene conversion, each subset of interest was analyzed using the HOMER software tool [18]. The software was configured to use the whole set of genes represented in the HumanMethylation450 architecture as a background. HOMER tested the genes in each subset of interest against 21 different databases, including the Gene Ontology (GO) Biological Process, Molecular Function and Cellular Component ontologies, as well as KEGG and Reactome pathway databases, among many others.

### Circular visualization and track smoothing

In order to plot the CpG and histone peak information on the circular genome-wide and example graphs, smoothing was applied to the data. CpG enrichment information for canonical and non-canonical hypermethylation was generated by partitioning the genome into intervals of 10kbp and assigning to each a score corresponding to the average coverage of the selected CpGs in the interval.

### Whole-genome bisulfite sequencing (WGBS) datasets

Supplementary data referenced in [42] was used as a validation dataset in glioblastoma. Previously processed data in the form of quantified methylation for each CpG measured in both strands of the genome was downloaded and filtered. Only methylation measures from CpGs having a total read count higher than 10 were retained.

The resulting dataset comprised only two samples (normal and tumoral), so a descriptive strategy was used to distinguish the different types of probes according to their methylation status. Hydroxymethylated probes were identified as those having a 5hmC measure higher than 0.1. Differentially methylated probes were defined as those having an absolute difference in their methylation values, between the control and tumor samples, higher than a given threshold (0.2 for 5mC and 0.1 for 5hmC).

The validation datasets may contain either one or two methylation measures for each CpG in the genome as they measure methylation in both strands. Strand-agnostic CpG regions representing the CpG dinucleotides with at least one measure were defined in order to compute the degree of intersection between the WGBS and methylation arrays results.

### TCGA expression datasets

In order to analyze changes in gene expression, samples of glioblastoma multiforme (GBM) were selected from among the data generated by the TCGA Research Network (http://cancergenome.nih.gov). Expression Level-3 pre-processed data was obtained for 572 GBM samples (10 controls and 562 tumors). The moderated t-test approach in the R/Bioconductor package *limma* was used to assess the differential expression status of each gene in the TCGA datasets. The normalized expression ratio in the TCGA datasets was used as the response variable, and the sample group (normal/tumoral) as the covariate of interest. No adjustment for possible confounders was performed in this case. An FDR threshold of 0.001 was used to correct for multiple hypotheses. No filtering on effect size was applied in this case.

### Data analysis workflow

All the necessary steps for upstream and downstream analyses were defined and implemented using the Snakemake tool [26], which helps data scientists to generate a reproducible and inherently parallel processing pipeline. Individual workflow tasks were implemented in R (version 3.2.2) and Python (version 3.4.3).

## Acknowledgments

We thank Ronnie Lendrum for editorial assistance. We also thank the Tumor Bank of the Hospital Virgen de la Salud (BioB-HVS, Toledo, Spain) for providing tumor samples. This work has been financially supported by: the Plan Nacional de I+D+I 2013-2016/FEDER (PI15/00892 to M.F.F. and A.F.F.); the ISCIII-Subdirección General de Evaluación y Fomento de la Investigación, and the Plan Nacional de I+D+I 2008-2011/FEDER (CP11/00131 to A.F.F.); IUOPA (to G.F.B. and M.S); the Fundación Científica de la AECC (to R.G.U.); the Fundación Ramón Areces (to M.F.F); FICYT (to E.G.T., M.G.G. and A.C.); and the Asturias Regional Government (GRUPIN14-052 to M.F.F.). Work in P.M. lab is supported by the European Research Council (CoG-2014-646903), the Spanish Ministry of Economy-Competitiveness (SAF-SAF2013-43065), the Obra Social La Caixa-Fundaciò Josep Carreras, and the Generalitat de Catalunya. P.M. is an investigator in the Spanish Cell Therapy cooperative network (TERCEL). The IUOPA is supported by the Obra Social Cajastur-Liberbank, Spain.

## Supplementary files

Supplementary Tables 1-12

Supplementary figures 1-4

**Figure 4-figure supplement 1.**
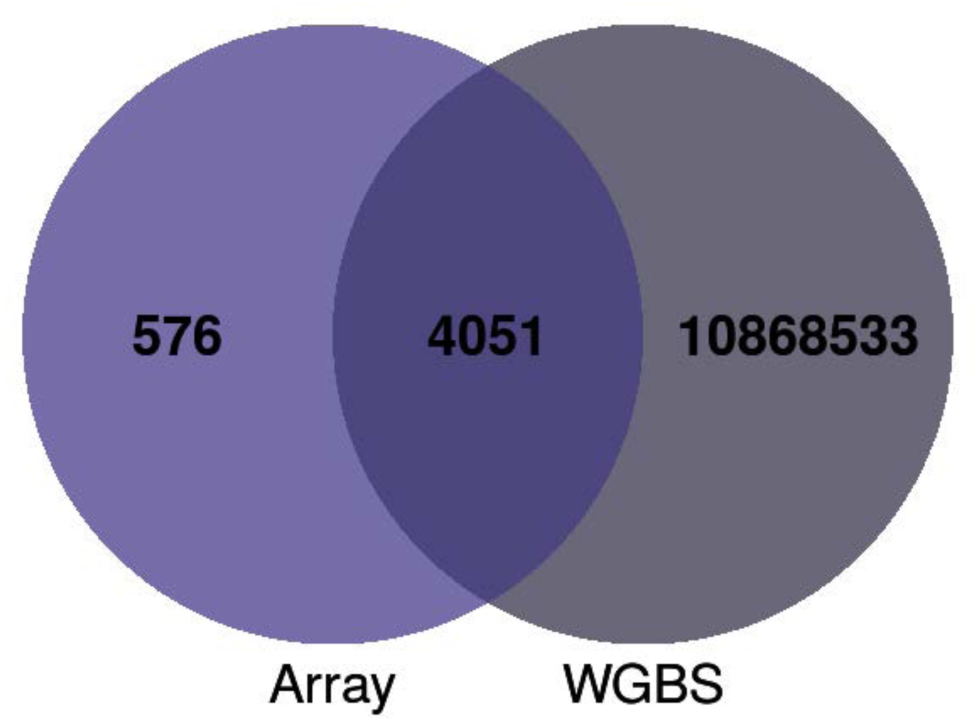
Venn diagram showing the overlap of hyper5mC-hypo5hmC sites in glioma obtained by methylation arrays and whole genome bisulfite sequencing (WGBS).

**Figure 5-figure supplement 1.**
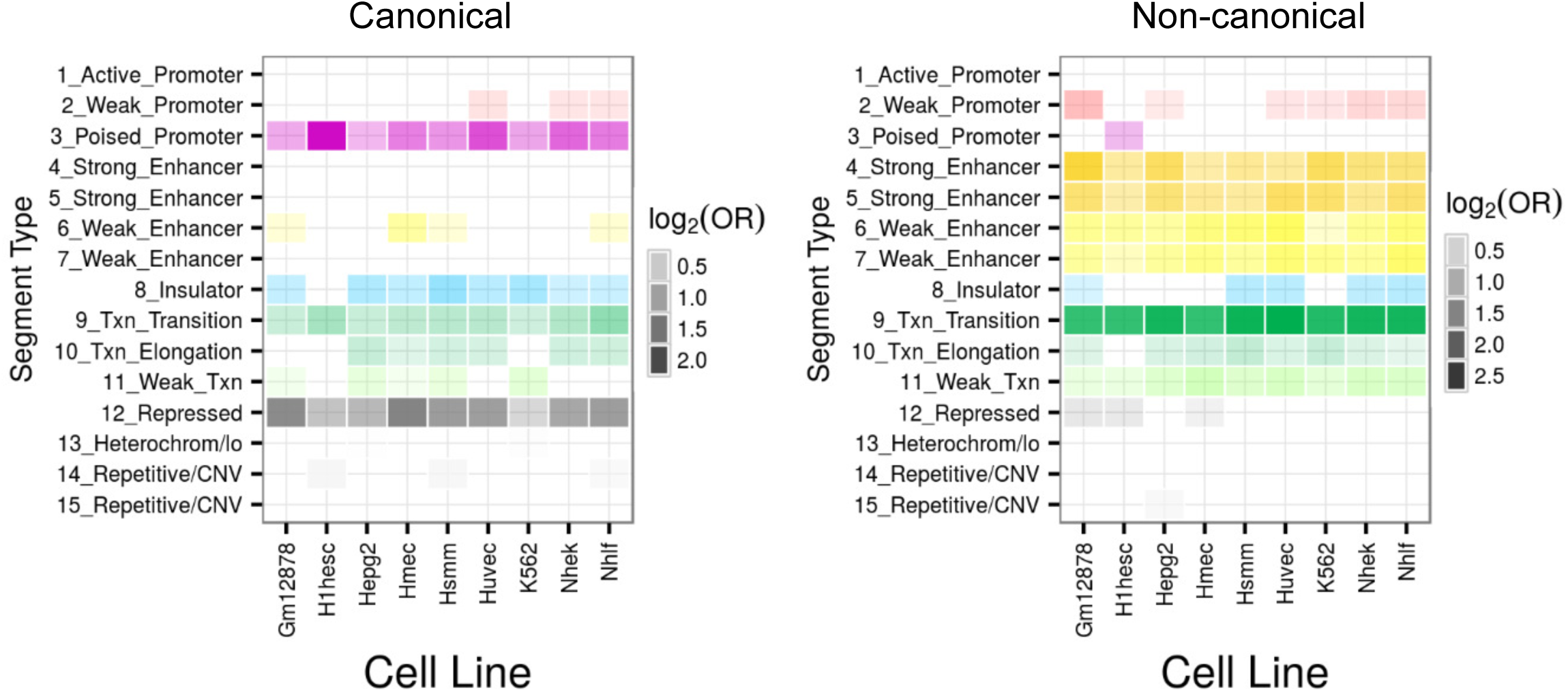
Canonical and non-canonical hypermethylation in glioma. Heatmaps showing significant enrichment of canonical (left panel) and non-canonical (right panel) hypermethylated CpG sites with fifteen “chromatin states” generated by a Hidden Markov Model (HMM). Colour codes indicate the significant enrichment based on log2 odds ratio (OR).

**Figure 6-figure supplement 1.**
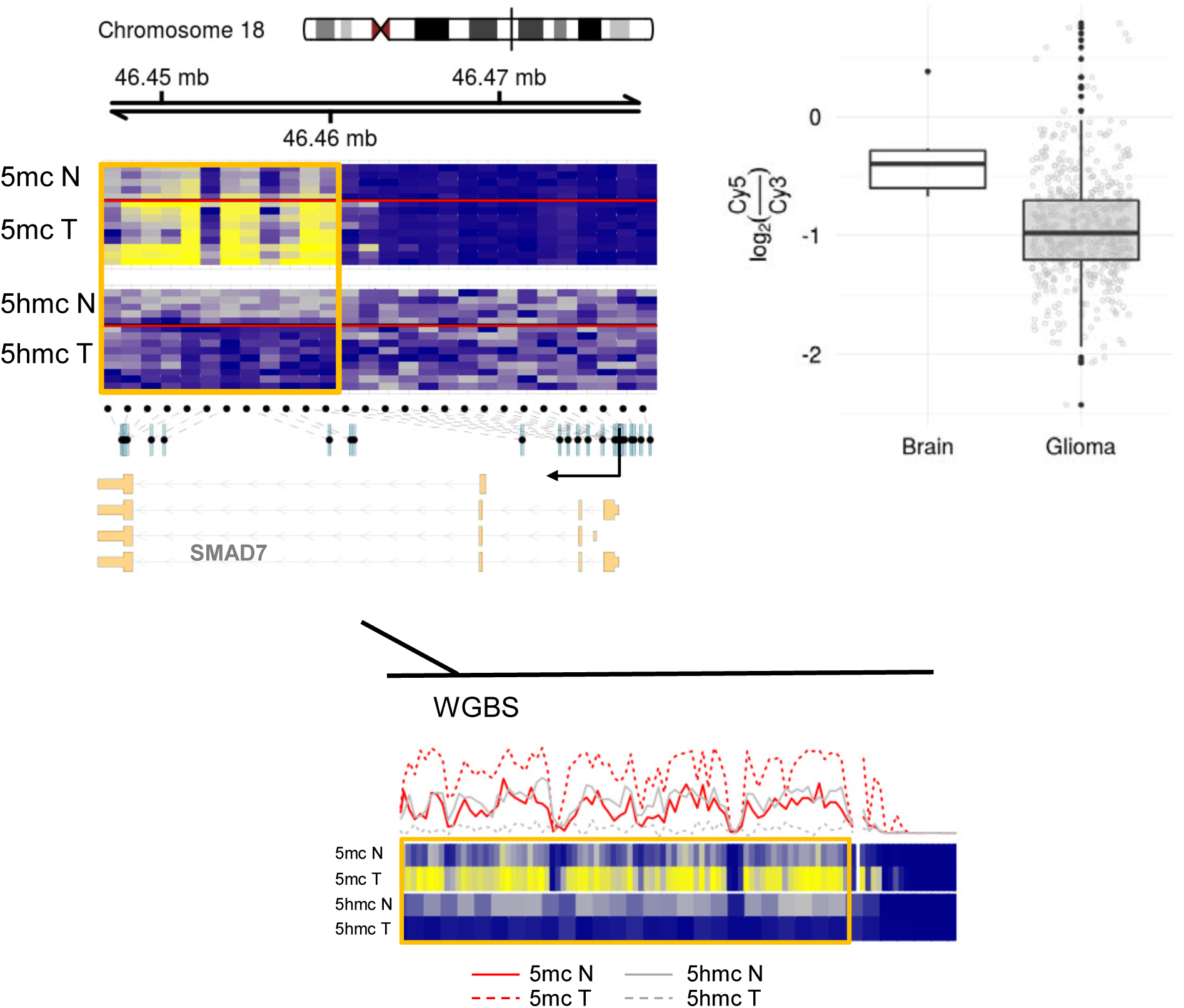
Functional role of non-canonical hypermethylation in glioma. Representative example of one gene (*SMAD7*) showing non-canonical hypermethylation in glioma (orange frame). Organization of the gene, location of CpGs included in the methylation array (black dots), and transcription start site (TSS) are shown below. 5mC hypermethylation (blue to yellow) and 5hmC loss (gray to blue) in gliomas are shown above. Lower panel shows the full genome bisulfite sequencing (WGBS) data including all the CpG sites in the same region. The associated change in gene expression is shown on the right.

**Figure 2-figure supplement 1.**
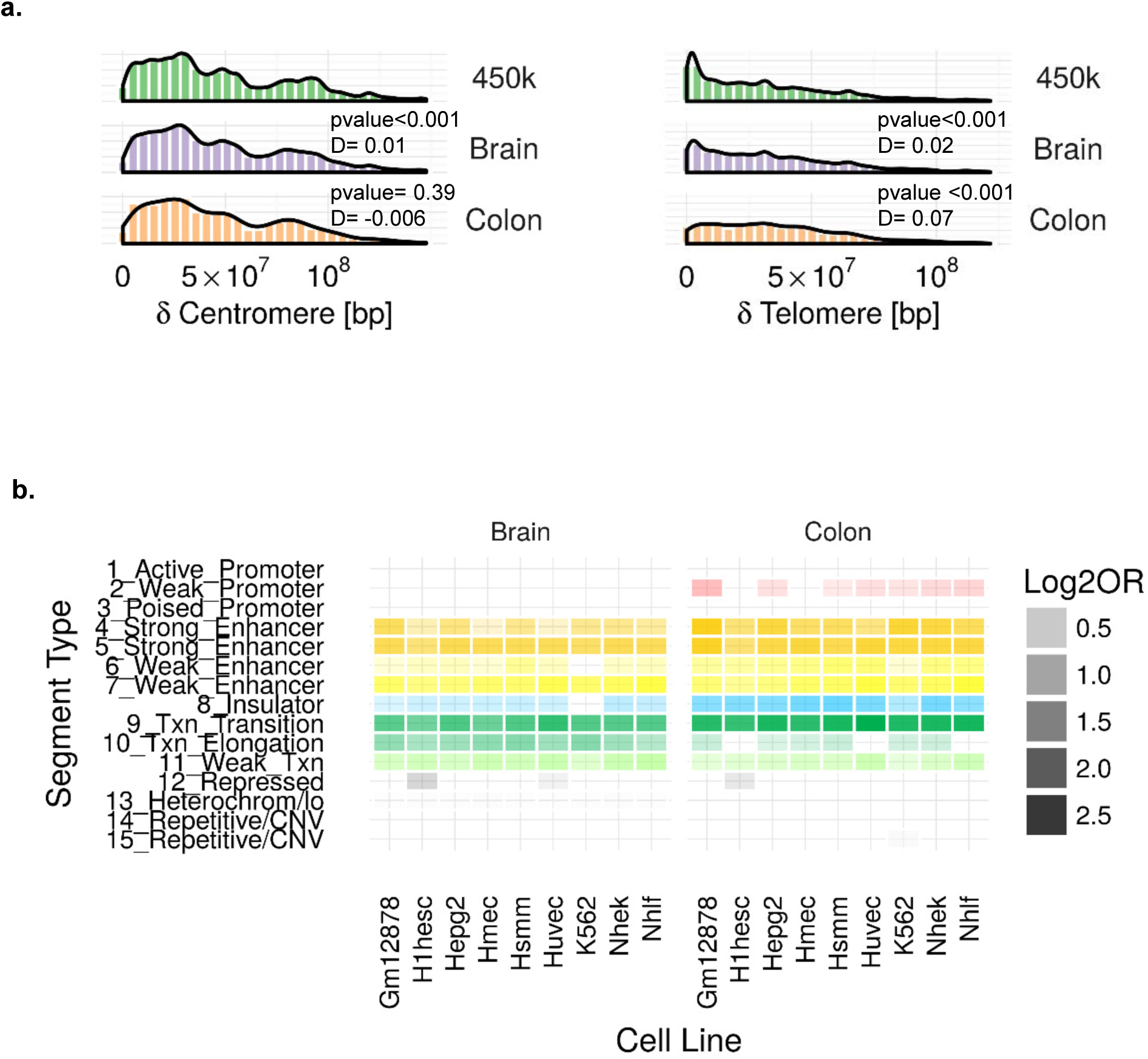
Characterization of DNA 5hmC in normal brain and colon samples. **(a)** Histograms and density plots showing the associations between 5hmC CpGs and distance to centromere (left) and telomeres (right). **(b)** Heatmaps showing significant enrichment of 5hmC CpG sites with fifteen “chromatin states” generated by a Hidden Markov Model (HMM). Colour codes indicate the significant enrichment based on log2 odds ratio (OR).

